# Enhanced transmissibility, infectivity and immune resistance of the SARS-CoV-2 Omicron XBB.1.5 variant

**DOI:** 10.1101/2023.01.16.524178

**Authors:** Keiya Uriu, Jumpei Ito, Jiri Zahradnik, Shigeru Fujita, Yusuke Kosugi, Gideon Schreiber, The Genotype to Phenotype Japan (G2P-Japan) Consortium, Kei Sato

## Abstract

In 2022, we have elucidated the characteristics of a variety of newly emerging SARS-CoV-2 Omicron subvariants. At the end of 2022, the XBB.1.5 variant, an descendant of XBB.1 that acquired the S:F486P substitution, emerged and is rapidly spreading in the USA and is the latest variant of concern. Although the features of XBB.1.5 was already reported by another group as a preprint, we think multiple and independent evaluations important, and these reports are crucial for sustained global health. In this study, our epidemic dynamics analysis revealed that the relative effective reproduction number (Re) of XBB.1.5 is more than 1.2-fold greater than that of the parental XBB.1, and XBB.1.5 is outcompeting BQ.1.1, the predominant lineage in the USA as of December 2022. Our data suggest that XBB.1.5 will rapidly spread worldwide in the near future. Yeast surface display assay and pseudovirus assay respectively showed that the ACE2 binding affinity and infectivity of XBB.1.5 is 4.3-fold and 3.3-fold higher than those of XBB.1, respectively. Moreover, neutralization assay revealed that XBB.1.5 is robustly resistant to BA.2 breakthrough infection sera (41-fold versus B.1.1, 20-fold versus BA.2) and BA.5 breakthrough infection sera (32-fold versus B.1.1, 9.5-fold versus BA.5), respectively. Because the immune resistance of XBB.1.5 is comparable to that of XBB.1, our results suggest that XBB.1.5 is the most successful XBB lineage as of January 2023 by acquiring the S:F486P substitution to augment ACE2 binding affinity without losing remarkable immune resistance, which leads to greater transmissibility.

## Main

In late 2022, the SARS-CoV-2 Omicron BQ.1 and XBB lineages, characterized by amino acid substitutions in the spike (S) proteins to increase viral fitness, have become predominant in the Western and Eastern Hemisphere, respectively.^1,2^ The BQ.1 lineages are descendants of BA.5, while the XBB lineage is the recombinant of two highly diversified BA.2 lineages.^2^

In 2022, we have elucidated the characteristics of a variety of newly emerging SARS-CoV-2 Omicron subvariants.^1-6^ At the end of 2022, the XBB.1.5 variant, an descendant of XBB.1 that acquired the S:F486P substitution, emerged and is rapidly spreading in the USA (**Figure 1A**), and is the latest variant of concern.^7^ Although the features of XBB.1.5 were reported by Yue et al.,^8^ a comprehensive understanding of the virological characteristics of newly emerging variants is needed for sustained global health. Our epidemic dynamics analysis (see **Appendix**) revealed that the relative effective reproduction number (R_e_) of XBB.1.5 is more than 1.2-fold greater than that of the parental XBB.1, and XBB.1.5 is outcompeting BQ.1.1, the predominant lineage in the USA as of December 2022 (**Figures 1A and 1B and Table S1**). Our data suggest that XBB.1.5 will rapidly spread worldwide in the near future (**Figure 1B**). We also found that a part of XBB.1.5 lost the deletion of Y at residue 144 in S (S:Y144del), which increases immune escape ability but decreases viral infectivity.^2^ However, the XBB.1.5 without S:Y144del (XBB.1.5+ins144Y) showed relatively lower R_e_ than the original XBB.1.5 (**Figure 1B**).

**Figure 1.**
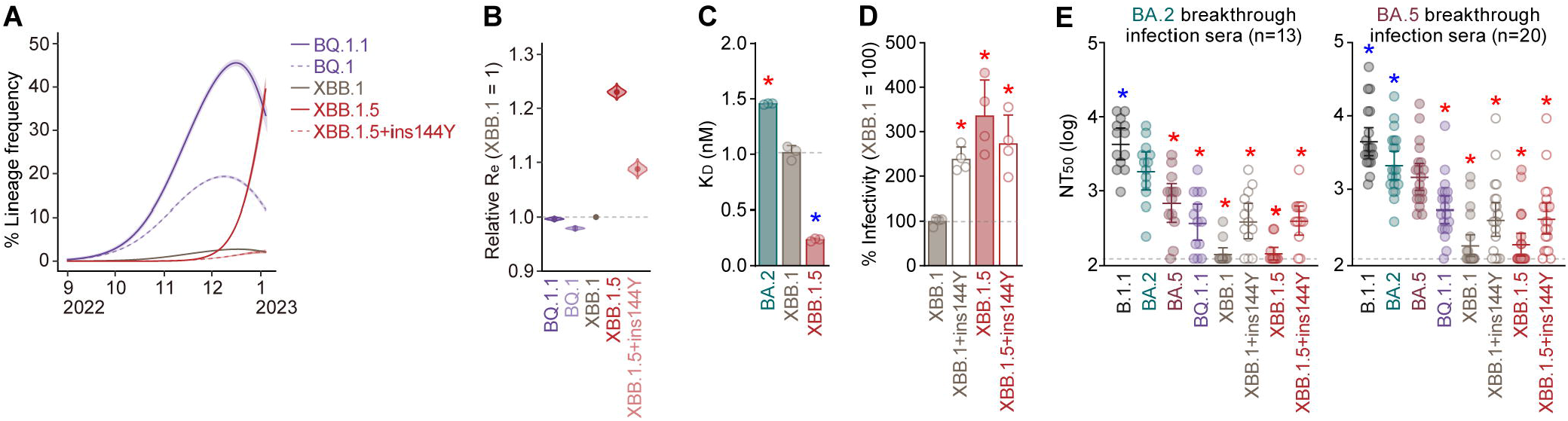
Virological features of Omicron XBB.1.5. **(A)** Estimated epidemic dynamics of representative viral lineages in the USA [posterior mean, line; 95% Bayesian confidence interval (CI), ribbon]. **(B)** Estimated relative R_e_ for each viral lineage. The R_e_ value of XBB.1 is set at 1. The posterior (violin), posterior mean (dot), and 95% CI (line) are shown. The raw data are summarized in **Table S1**. **(C)** Binding affinity of the RBD of SARS-CoV-2 S protein to ACE2 by yeast surface display. The K_D_ value indicating the binding affinity of the RBD of the SARS-CoV-2 S protein to soluble ACE2 when expressed on yeast is shown. **(D)** Pseudovirus assay. HOS-ACE2-TMPRSS2 cells were infected with pseudoviruses bearing each S protein. The amount of input virus was normalized based on the amount of HIV-1 p24 capsid protein. The percent infectivity compared to that of the virus pseudotyped with the XBB.1 S protein are shown. **(E)** Neutralization assay. Assays were performed with pseudoviruses harboring the S proteins of B.1.1, BA.2, BA.5, BQ.1.1, XBB.1, XBB.1+ins144Y, XBB.1.5, XBB.1.5+ins144Y. Convalescent sera from fully vaccinated individuals who had been infected with BA.2 after full vaccination (9 2-dose vaccinated and 4 3-dose vaccinated. 13 donors in total) (left) and those who had been infected with BA.5 after full vaccination (2 2-dose vaccinated donors, 17 3-dose vaccinated donors and 1 4-dose vaccinated donors. 20 donors in total) (right) were used. The horizontal dashed line indicates the detection limit (120-fold). In **C** and **D**, assays were performed in triplicate (**C**) or quadruplicate (**D**). The presented data are expressed as the average ± SD. Statistically significant differences (*, *P* < 0.05) versus XBB.1 were determined by two-sided Student’s *t* tests. Red and blue asterisks, respectively, indicate increased and decreased values. The horizontal dashed line indicates the value of XBB.1. In **E**, each dot indicates the result of an individual replicate. Assays for each serum sample were performed in triplicate to determine the 50% neutralization titer (NT_50_). Each dot represents one NT_50_ value, and the geometric mean and 95% CI are shown. Statistically significant differences (*, *P* < 0.05) versus XBB.1 were determined by two-sided Wilcoxon signed-rank tests and indicated with asterisks. Red and blue asterisks, respectively, indicate decreased and increased NT_50_ values. Information on the convalescent donors is summarized in **Table S2**.

We next investigated the virological features of XBB.1.5. Yeast surface display assay showed that the K_D_ value of XBB.1.5 S receptor-binding domain (RBD) to human ACE2 receptor is significantly (4.3-fold) lower than that of XBB.1 S RBD (**Figure 1C**). Experiments using pseudoviruses also showed approximately 3-fold increased infectivity of XBB.1.5 compared to XBB.1 (**Figure 1D**). These results suggest that XBB.1.5 exhibits remarkably strong affinity to human ACE2, which is attributed to the F486P substitution. On the other hand, the 144Y insertion mutation increased the XBB.1 infectivity but did not that of XBB.1.5 infectivity (**Figure 1D**).

Finally, neutralization assay revealed that XBB.1.5 is robustly (41-fold versus B.1.1, 20-fold versus BA.2) resistant to BA.2 breakthrough infection sera (**Figure 1E**). XBB.1.5 is also severely (32-fold versus B.1.1, 9.5-fold versus BA.5) resistant to BA.5 breakthrough infection sera (**Figure 1E**). The 144Y insertion significantly increased the sensitivity to both BA.2 and BA.5 breakthrough infection sera (**Figure 1E**).

In sum, our results suggest that XBB.1.5 is the most successful XBB lineage as of January 2023 by acquiring the S:F486P substitution to augment ACE2 binding affinity without losing remarkable immune resistance, which leads to greater transmissibility.

## Grants

Supported in part by AMED SCARDA Japan Initiative for World-leading Vaccine Research and Development Centers “UTOPIA” (JP223fa627001, to Kei Sato), AMED SCARDA Program on R&D of new generation vaccine including new modality application (JP223fa727002, to Kei Sato); AMED Research Program on Emerging and Re-emerging Infectious Diseases (JP22fk0108146, to Kei Sato; JP21fk0108494 to G2P-Japan Consortium and Kei Sato; JP21fk0108425, to Kei Sato; JP21fk0108432, to Kei Sato); AMED Research Program on HIV/AIDS (JP22fk0410039, to Kei Sato); JST PRESTO (JPMJPR22R1, to Jumpei Ito); JST CREST (JPMJCR20H4, to Kei Sato); JSPS KAKENHI Grant-in-Aid for Early-Career Scientists (20K15767, Jumpei Ito); JSPS Core-to-Core Program (A. Advanced Research Networks) (JPJSCCA20190008, Kei Sato); JSPS Research Fellow DC2 (22J11578, to Keiya Uriu); The Tokyo Biochemical Research Foundation (to Kei Sato); and the project of National Institute of Virology and Bacteriology, Programme EXCELES, funded by the European Union, Next Generation EU (LX22NPO5103, to Jiri Zahradnik).

