## Supplementary Appendix for "Enhanced transmissibility, infectivity and immune resistance of the SARS-CoV-2 Omicron XBB.1.5 variant"

**Table of Contents**

**Contents** Page


| | |
|---|---|
| **Materials and Methods** | 2-4 |
| Ethics statement | |
| Human serum collection | |
| Epidemic dynamic analysis | |
| Plasmid construction | |
| Yeast surface display | |
| Cell culture | |
| Neutralization assay | |
| Data availability | |
| **Table S1.** Information on estimated relative $R_e$ value for each viral lineage | 5-6 |
| **Table S2.** Human sera used in this study | 7 |
| **Table S3.** Primers used in this study | 8 |
| **Consortia** | 9 |
| **Acknowledgments** | 10 |
| **Supplemental References** | 11-12 |

### Materials and Methods

#### Ethics statement

All protocols involving specimens from human subjects recruited at Interpark Kuramochi Clinic was reviewed and approved by the Institutional Review Board of Interpark Kuramochi Clinic (approval ID: G2021-004). All human subjects provided written informed consent. All protocols for the use of human specimens were reviewed and approved by the Institutional Review Boards of The Institute of Medical Science, The University of Tokyo (approval IDs: 2021-1-0416 and 2021-18-0617).

#### Human serum collection

Convalescent sera were collected from fully vaccinated individuals who had been infected with BA.2 (9 2-dose vaccinated and 4 3-dose vaccinated; 11–61 days after testing. n=13 in total; average age: 45 years, range: 24–82 years, 62% male) (**Figure 1E**), and fully vaccinated individuals who had been infected with BA.5 (2 2-dose vaccinated, 17 3-dose vaccinated and 1 4-dose vaccinated; 10–23 days after testing. n=20 in total; average age: 51 years, range: 25–73 years, 45% male) (**Figure 1E**). The SARS-CoV-2 variants were identified as previously described.[1-3] Sera were inactivated at 56°C for 30 minutes and stored at –80°C until use. The details of the convalescent sera are summarized in **Table S2**.

#### Epidemics dynamics analysis

We modeled the epidemic dynamics of viral lineages in the USA based on the viral genomic surveillance data deposited in the GISAID database (https://www.gisaid.org/; downloaded on January 9th, 2023). In the present study, we analyzed the data from September 1, 2022. We excluded the sequence records with the following features: i) a lack of collection date information; ii) sampling in animals other than humans; iii) sampling by quarantine; iv) without the PANGO lineage information; and v) having >2% undetermined (N) nucleotide sequences. Sublineages of BQ.1.1 are summarized as BQ.1.1, and other BQ.1 sublineages are summarized as BQ.1. In addition, according to the presence or absence of S:Y144del, we classified XBB.1.5 into two groups, XBB.1.5 and XBB.1.5+ins144Y (XBB.1.5 without S:Y144del). We removed XBB from the analysis since we found that various XBB sublineages are contaminated into this category. In the downstream analysis, we only used sequences for PANGO lineages with >500 sequences in the dataset. We counted the daily frequency of each viral lineage. Subsequently, epidemic dynamics and relative $R_e$ value for each viral lineage were estimated according to the Bayesian multinomial logistic model, described in our previous study.[2] Briefly, we estimated the logistic slope parameter $\beta_l$ for each viral lineage using the model and then calculated relative $R_e$ for each lineage $r_l$ as $r_l = exp(\gamma\beta_l)$ where $\gamma$ is the average viral generation time (2.1 days) (http://sonorouschocolate.com/covid19/index.php?title=Estimating_Generation_Time_Of_Omicr on). For parameter estimation, the intercept and slope parameters of XBB.1 were fixed at 0. Consequently, the relative $R_e$ of XBB.1 was fixed at 1, and those of the other lineages were estimated relative to that of XBB.1. Parameter estimation was performed via the MCMC approach implemented in CmdStan v2.31.0 (https://mc-stan.org) with CmdStanr v0.5.3 (https://mc-stan.org/cmdstanr/). Four independent MCMC chains were run with 1,000 and 4,000 steps in the warmup and sampling iterations, respectively. We confirmed that all estimated parameters showed <1.01 R-hat convergence diagnostic values and >200 effective sampling size values, indicating that the MCMC runs were successfully convergent. Information on the estimated parameters is summarized in **Table S1**. In **Figure 1A**, results for BQ.1.1, BQ.1, XBB.1, XBB.1.5, and XBB.1.5+ins144Y are shown.

### Plasmid construction

Plasmids expressing the SARS-CoV-2 spike proteins of the parental D614G (B.1.1), Omicron BA.2, BA.5, BQ.1.1 and XBB.1 were prepared in our previous studies.[1,2,4-8] Plasmids expressing the spike protein of XBB.1.5 and its derivative were generated by site-directed overlap extension PCR using pC-SARS2-S XBB.1[8] as the template and the primers listed in **Table S3**. The resulting PCR fragment was subcloned into the KpnI-NotI site of the pCAGGS vector[9] using In-Fusion® HD Cloning Kit (Takara, Cat# Z9650N). Nucleotide sequences were determined by DNA sequencing services (Eurofins), and the sequence data were analyzed by Sequencher v5.1 software (Gene Codes Corporation).

### Yeast surface display

Yeast surface display binding analyses for the spike receptor-binding domains of BA.2, XBB and XBB.1.5 (residues 333–527) were performed as previously described.[1-4,7,8,10-12] Cerevisiae EBY100 yeasts and pJYDC1 plasmids with RBD genes cloned between the NdeI and BamHI sites (Addgene, Cat# 162458) as previously described.[10,11] Transformed yeasts were grown in SDCAA media (220 rpm, 30°C) and expressed overnight in expression media supplemented with bilirubin (10 nM DMSO solubilized, Sigma-Aldrich, Cat# 14370) after inoculation to OD600 0.7–1.0 (220 rpm, 20°C).[10,11] Aliquots of expressed yeast cells (100 μl) were washed in ice-cold PBSB buffer (PBS with 1 mg/ml BSA) and incubated in a series of CF®640R succinimidyl ester labeled (Biotium, Cat# 92108) ACE2 peptidase domain (residues 18–740) concentrations, PBSB buffer and 1 nM bilirubin for 8 to 12 hours. After incubation, the unbound fraction was washed by ice-cold PBSB buffer and yeasts were transferred into a 96-well plate (Thermo Fisher Scientific, Cat# 268200) and 30,000 events in gated population were automatically acquired by a CytoFLEX S Flow Cytometer (Beckman Coulter, USA, Cat#. N0-V4-B2-Y4) with FITC channel data for eUnaG2 fluorescence (Abs/Em maxima 498/527 nm) and APC channel for CF640 fluorescence (Abs/Em maxima 642/662 nm) setting. Gating, analysis and fitting with nonlinear least-squares regression using Python v3.7 protocols were described previously.[1-4,7,8,10-12]

**Cell culture**

HEK293T cells (a human embryonic kidney cell line; ATCC CRL-3216) and HOS-ACE2/TMPRSS2 cells (kindly provided by Dr. Kenzo Tokunaga),[13,14] a derivative of HOS cells (a human osteosarcoma cell line; ATCC CRL-1543) stably expressing human ACE2 and TMPRSS2, were maintained in Dulbecco's modified Eagle's medium (DMEM) (high glucose) (Wako, Cat# 044-29765) containing 10% fetal bovine serum (FBS) (Sigma-Aldrich Cat# 172012-500ML), 100 units penicillin and 100 ug/ml streptomycin (PS) (Sigma-Aldrich, Cat# P4333-100ML).

**Neutralization assay**

Pseudoviruses were prepared as previously described.[1-3,5-8,12,13,15-18] Briefly, lentivirus (HIV-1)-based, luciferase-expressing reporter viruses were pseudotyped with the SARS-CoV-2 spikes. HEK293T cells ($1 \times 10^6$ cells) were cotransfected with 1 μg psPAX2-IN/HiBiT,[19] 1 μg pWPI-Luc2,[19] and 500 ng plasmids expressing parental S or its derivatives using PEI Max (Polysciences, Cat# 24765-1) according to the manufacturer's protocol. Two days post transfection, the culture supernatants were harvested and centrifuged. The pseudoviruses were stored at –80°C until use. Neutralization assays were performed as previously described.[1-3,5-8,12,13,15-18] Briefly, the SARS-CoV-2 spike pseudoviruses (counting ~20,000 relative light units) were incubated with serially diluted (120-fold to 87,480-fold dilution at the final concentration) heat-inactivated sera at 37°C for 1 hour. Pseudoviruses without sera were included as controls. Then, an 40 μl mixture of pseudovirus and serum was added to HOS-ACE2/TMPRSS2 cells (10,000 cells/50 μl) in a 96-well white plate. Two days post infection, the infected cells were lysed with a Bright-Glo luciferase assay system (Promega, Cat# E2620), and the luminescent signal was measured using a GloMax explorer multimode microplate reader 3500 (Promega). The assay of each serum sample was performed in triplicate, and the 50% neutralization titer was calculated using Prism 9 (GraphPad Software).

**Data availability**

Dataset used in the epidemic dynamics analysis in this study is available from the GISAID database (https://www.gisaid.org; EPI_SET_230113qo). The GISAID supplemental tables for EPI_SET_230113qo is available in the GitHub repository (https://github.com/TheSatoLab/XBB.1.5_short).

**Table S1. Information on estimated relative Re value for each viral lineage**

| PANGO lineage | Posterior mean | Posterior 2.5 percentile | Posterior 97.5 percentile | R-hat value | Effective sampling size (ESS_bulk) | Effective sampling size (ess_tail) |
|---|---|---|---|---|---|---|
| XBB.1.5 | 1.234 | 1.222 | 1.246 | 1.002 | 1952.4 | 4951.0 |
| XBB.1.5+ins144Y | 1.089 | 1.075 | 1.102 | 1.001 | 2894.9 | 7866.7 |
| XBB.2 | 1.002 | 0.993 | 1.011 | 1.003 | 1729.2 | 4963.0 |
| BQ.1.1 | 0.997 | 0.992 | 1.001 | 1.009 | 390.3 | 854.0 |
| CK.1 | 0.996 | 0.987 | 1.005 | 1.003 | 1624.0 | 4448.7 |
| CQ.2 | 0.991 | 0.981 | 1.001 | 1.002 | 1762.0 | 5079.7 |
| BQ.1 | 0.979 | 0.975 | 0.983 | 1.009 | 395.6 | 943.6 |
| BA.5.2.35 | 0.966 | 0.958 | 0.974 | 1.002 | 1484.2 | 4403.4 |
| BA.2.75 | 0.941 | 0.936 | 0.945 | 1.008 | 436.2 | 1056.3 |
| BA.5.1.27 | 0.939 | 0.934 | 0.945 | 1.004 | 790.9 | 2159.1 |
| BA.5.2.6 | 0.938 | 0.933 | 0.943 | 1.006 | 577.2 | 1502.3 |
| BF.14 | 0.933 | 0.926 | 0.940 | 1.002 | 1025.7 | 2773.6 |
| BF.7.4.1 | 0.933 | 0.927 | 0.938 | 1.004 | 710.3 | 2026.7 |
| CM.2 | 0.930 | 0.923 | 0.937 | 1.003 | 1191.1 | 3681.0 |
| BA.5.2.34 | 0.927 | 0.921 | 0.932 | 1.005 | 673.0 | 1735.0 |
| BF.7 | 0.920 | 0.916 | 0.924 | 1.008 | 435.9 | 1043.3 |
| BF.11 | 0.917 | 0.912 | 0.922 | 1.006 | 599.0 | 1661.9 |
| BE.1.1.1 | 0.916 | 0.909 | 0.922 | 1.003 | 933.2 | 2275.5 |
| BA.5.2.23 | 0.915 | 0.909 | 0.921 | 1.004 | 838.2 | 2615.1 |
| BF.7.4 | 0.915 | 0.910 | 0.920 | 1.006 | 663.8 | 1894.7 |
| BA.5.1.18 | 0.898 | 0.893 | 0.904 | 1.005 | 714.5 | 1916.6 |
| BA.5.9 | 0.893 | 0.886 | 0.900 | 1.003 | 1134.7 | 3412.8 |
| BF.13 | 0.885 | 0.879 | 0.891 | 1.004 | 831.7 | 2318.6 |
| BA.5.1.5 | 0.884 | 0.879 | 0.889 | 1.005 | 734.6 | 2080.0 |
| BE.1.1 | 0.884 | 0.879 | 0.888 | 1.006 | 537.7 | 1402.8 |
| BF.26 | 0.881 | 0.876 | 0.885 | 1.007 | 509.9 | 1284.5 |
| BA.5.5.1 | 0.871 | 0.866 | 0.877 | 1.004 | 868.0 | 2493.4 |
| BA.5.2.3 | 0.869 | 0.862 | 0.875 | 1.002 | 1198.5 | 3518.6 |
| BA.5.2.20 | 0.866 | 0.861 | 0.871 | 1.005 | 667.4 | 2045.0 |
| BA.5.1.22 | 0.865 | 0.860 | 0.870 | 1.005 | 718.1 | 2217.6 |
| BA.5.2.31 | 0.865 | 0.858 | 0.871 | 1.003 | 1054.5 | 2983.0 |
| BE.1.4 | 0.862 | 0.856 | 0.869 | 1.003 | 1173.4 | 3736.9 |
| BA.5.2 | 0.862 | 0.858 | 0.866 | 1.009 | 393.4 | 862.8 |
| BA.5.1.10 | 0.861 | 0.856 | 0.866 | 1.006 | 650.1 | 1720.5 |
| BA.5.1.6 | 0.861 | 0.855 | 0.868 | 1.004 | 1108.2 | 2816.6 |
| BA.5 | 0.861 | 0.857 | 0.865 | 1.007 | 505.0 | 1329.1 |
| BA.5.2.21 | 0.861 | 0.856 | 0.865 | 1.005 | 595.0 | 1765.2 |
| BF.10 | 0.859 | 0.855 | 0.863 | 1.008 | 461.5 | 1127.6 |
| BA.5.1.3 | 0.858 | 0.851 | 0.865 | 1.002 | 1302.3 | 3986.9 |
| BF.5 | 0.857 | 0.853 | 0.862 | 1.007 | 527.5 | 1417.6 |
| BA.5.2.9 | 0.856 | 0.852 | 0.860 | 1.007 | 494.7 | 1199.0 |
| BA.4.6.5 | 0.855 | 0.850 | 0.861 | 1.004 | 808.8 | 2293.9 |

(Table S1, continued)

| | | | | | | |
|---|---|---|---|---|---|---|
| BA.5.1 | 0.855 | 0.851 | 0.859 | 1.008 | 415.0 | 957.0 |
| BA.5.1.23 | 0.855 | 0.849 | 0.860 | 1.004 | 786.8 | 1947.4 |
| BA.4.6 | 0.854 | 0.850 | 0.858 | 1.008 | 398.5 | 887.0 |
| BA.5.1.25 | 0.851 | 0.845 | 0.858 | 1.003 | 1203.5 | 3445.9 |
| BA.5.2.1 | 0.850 | 0.847 | 0.854 | 1.009 | 383.5 | 811.7 |
| BE.1 | 0.849 | 0.843 | 0.854 | 1.005 | 745.1 | 2051.5 |
| BA.5.1.30 | 0.847 | 0.841 | 0.853 | 1.004 | 999.1 | 2768.6 |
| BA.5.1.2 | 0.847 | 0.840 | 0.854 | 1.003 | 1193.7 | 3556.3 |
| BA.5.5 | 0.844 | 0.840 | 0.848 | 1.007 | 505.6 | 1218.6 |
| BA.5.2.22 | 0.843 | 0.836 | 0.850 | 1.003 | 1291.3 | 4064.0 |
| BF.27 | 0.837 | 0.831 | 0.843 | 1.003 | 909.9 | 2698.9 |
| BF.21 | 0.836 | 0.829 | 0.842 | 1.003 | 1081.6 | 3131.9 |
| BA.5.1.1 | 0.831 | 0.825 | 0.836 | 1.005 | 846.8 | 2285.5 |
| BF.8 | 0.830 | 0.823 | 0.837 | 1.002 | 1350.1 | 3848.4 |
| BA.5.6 | 0.819 | 0.815 | 0.824 | 1.007 | 577.2 | 1479.4 |
| BE.3 | 0.817 | 0.810 | 0.823 | 1.002 | 1237.1 | 3884.6 |
| BA.4.1 | 0.800 | 0.793 | 0.806 | 1.002 | 1496.2 | 4352.3 |

The Re value of XBB.1 is set at 1.

**Table S2. Human sera used in this study**

| SARS-CoV-2 variant infected | Donor ID | Sex | Age | Date of test (YYYY/MM/DD) | Date of sampling (YYYY/MM/DD) | Prior infection? | Prior vaccination? | Vaccine | Date of 1st vaccination (YYYY/MM/DD) | Date of 2nd vaccination (YYYY/MM/DD) | Date of 3rd vaccination (YYYY/MM/DD) | Date of 4rd vaccination (YYYY/MM/DD) |
|---|---|---|---|---|---|---|---|---|---|---|---|---|
| BA.2 | P378 | Male | 43 | 2022/03/28 | 2022/04/10 | No | Yes | BNT162b2 | 2021/10/10 | 2021/10/31 | | |
| BA.2 | P398 | Male | 48 | 2022/04/13 | 2022/04/30 | No | Yes | BNT162b2 | 2021/09/18 | 2021/10/09 | 2022/04/09 | |
| BA.2 | P407 | Male | 29 | 2022/05/01 | 2022/05/12 | No | Yes | mRNA-1273 | 2021/09/13 | 2021/10/11 | | |
| BA.2 | P401 | Male | 35 | 2022/04/22 | 2022/05/05 | No | Yes | BNT162b2 | 2021/09/09 | 2021/09/30 | | |
| BA.2 | P412 | Female | 82 | 2022/05/04 | 2022/05/26 | No | Yes | BNT162b2 | 2021/06/11 | 2021/07/09 | | |
| BA.2 | 6449 | Male | 43 | 2022/04/03 | 2022/04/23 | No | Yes | BNT162b2 | 2021/08/13 | 2021/09/11 | | |
| BA.2 | 6355 | Male | 50 | 2022/04/02 | 2022/04/20 | No | Yes | Not applicable | 2021/04/28 | 2021/05/19 | 2022/01/19 | |
| BA.2 | 6547 | Male | 54 | 2022/04/06 | 2022/04/22 | No | Yes | BNT162b2 | 2021/08/25 | 2021/09/15 | | |
| BA.2 | 7951 | Female | 71 | 2022/04/25 | 2022/05/12 | No | Yes | Mix | 2021/06/20 (BNT162b2) | 2021/07/16 (BNT162b2) | 2022/02/16 (mRNA-1273） | |
| BA.2 | 8645 | Female | 41 | 2022/05/07 | 2022/05/20 | No | Yes | BNT162b2 | 2021/05/23 | 2021/06/13 | 2022/01/20 | |
| BA.2 | 8682 | Female | 25 | 2022/05/08 | 2022/05/24 | No | Yes | BNT162b2 | 2021/09/03 | 2021/09/27 | | |
| BA.2 | 5949 | Male | 24 | 2022/03/22 | 2022/05/22 | No | Yes | mRNA-1273 | 2021/08/05 | 2021/09/02 | | |
| BA.2 | 8796 | Female | 34 | 2022/05/10 | 2022/06/05 | No | Yes | BNT162b2 | 2021/09/26 | 2021/10/17 | | |
| BA.5 | P427 | Female | 49 | 2022/07/06 | 2022/07/25 | No | Yes | Not applicable | 2021/07/30 | 2021/08/25 | 2022/03/18 | |
| BA.5 | P440 | Male | 25 | 2022/07/24 | 2022/08/07 | No | Yes | BNT162b2 | 2021/11/24 | 2021/12/15 | | |
| BA.5 | P439 | Female | 73 | 2022/07/23 | 2022/08/08 | No | Yes | BNT162b2 | 2021/06/19 | 2021/07/20 | 2022/02/04 (mRNA-1273) | |
| BA.5 | P451 | Female | 55 | 2022/07/29 | 2022/8/12 | No | Yes | BNT162b2 | 2021/04/26 | 2021/05/20 | 2022/01/18 | |
| BA.5 | P456 | Male | 44 | 2022/08/04 | 2022/08/14 | No | Yes | BNT162b2 | 2021/08/11 | 2021/09/01 | 2022/03/13 (mRNA-1273) | |
| BA.5 | P455 | Male | 29 | 2022/08/03 | 2022/08/17 | No | Yes | mRNA-1273 | 2021/07/26 | 2021/08/27 | 2022/04/18 | |
| BA.5 | P464 | Male | 63 | 2022/08/08 | 2022/08/19 | No | Yes | BNT162b2 | 2021/08/08 | 2021/08/29 | 2022/04/07 (mRNA-1273) | |
| BA.5 | 9341 | Male | 56 | 2022/06/12 | 2022/06/30 | No | Yes | BNT162b2 | 2021/08/10 | 2021/08/31 | 2022/03/18 | |
| BA.5 | 9584 | Male | 55 | 2022/07/08 | 2022/07/25 | No | Yes | BNT162b2 | 2021/07/14 | 2021/08/05 | 2022/03/22 | |
| BA.5 | 11318 | Female | 51 | 2022/07/24 | 2022/08/05 | No | Yes | BNT162b2 | 2021/09/01 | 2021/09/22 | 2022/05/19 (mRNA-1273) | |
| BA.5 | 23S-08 | Male | 25 | 2022/07/23 | 2022/08/08 | No | Yes | BNT162b2 | 2021/04/27 | 2021/05/18 | 2022/01/11 | |
| BA.5 | 11597 | Female | 41 | 2022/07/26 | 2022/08/08 | No | Yes | BNT162b2 | 2021/04/30 | 2021/05/21 | 2022/01/11 | |
| BA.5 | 10978 | Female | 46 | 2022/07/22 | 2022/08/11 | No | Yes | BNT162b2 | 2021/08/27 | 2021/09/17 | 2022/05/15 | |
| BA.5 | 10826 | Male | 63 | 2022/07/21 | 2022/08/11 | No | Yes | BNT162b2 | 2021/07/27 | 2021/08/17 | 2022/03/04 | 2022/8/9 (mRNA-1273) |
| BA.5 | 11079 | Female | 65 | 2022/07/23 | 2022/08/11 | No | Yes | mRNA-1273 | 2021/07/08 | 2021/08/05 | 2022/03/17 | |
| BA.5 | 14847 | Female | 70 | 2022/08/13 | 2022/08/25 | No | Yes | BNT162b2 | 2021/07/13 | 2021/08/20 | 2022/03/08 (mRNA-1273) | |
| BA.5 | 13180 | Female | 63 | 2022/08/04 | 2022/08/25 | No | Yes | BNT162b2 | 2021/07/16 | 2021/08/06 | 2022/03/08 (mRNA-1273) | |
| BA.5 | 12912 | Male | 64 | 2022/08/02 | 2022/08/25 | No | Yes | mRNA-1273 | 2021/09/02 | 2021/09/30 | 2021/04/01 | |
| BA.5 | 14956 | Female | 33 | 2022/08/13 | 2022/08/28 | No | Yes | BNT162b2 | 2021/09/06 | 2021/10/07 | | |
| BA.5 | 15707 | Female | 52 | 2022/08/16 | 2022/08/29 | No | Yes | BNT162b2 | 2021/08/07 | 2021/08/28 | 2022/04/03 | |

**Table S3. Primers used in this study**

| Primer name | Primer sequence (5'-to-3') | Use |
|---|---|---|
| Omicron universal Fw | cactatagggcgaattgggtaccatgtttgtgttcctggt | Preparation of S expression plasmid |
| BA2 Rv | agctccaccgcggtggcggccgctcaggtgtagtgcagtttca | Preparation of S expression plasmid |
| pC-S_XBB_S486P_Fwd | aatggagtggccggcCCCaactgttacAGCcca | Preparation of S expression plasmid |
| pC-S_XBB_S486P_Rev | tggGCTgtaacagttGGGgccggccactccatt | Preparation of S expression plasmid |
| pC-S_BA.2_F486P_Fwd | aatggagtggccggcCCCaactgttactttcca | Preparation of S expression plasmid |
| pC-S_BA.2_F486P_Rev | tggaaagtaacagttGGGgccggccactccatt | Preparation of S expression plasmid |
| pC-S_XBB_insY144_Fwd | ccattcctgGACgtcTACtacCAGaagaacaac | Preparation of S expression plasmid |
| pC-S_XBB_insY144_Rev | gttgttcttCTGgtaGTAgacGTCcaggaatgg | Preparation of S expression plasmid |
| XBB_F486P_F | CAGGCCGGTAACAAACCTTGTAATGGTGTTGCAGGTCC AAATTGTTACTCTCCTTTACAATCATATGGTTTCC | Preparation of S RBD expression plasmid |

### Consortia

#### The Genotype to Phenotype Japan (G2P-Japan) Consortium

**The Institute of Medical Science, The University of Tokyo, Japan**
Izumi Kimura, Naoko Misawa, Arnon Plianchaisuk, Lin Pan, Mai Suganami, Mika Chiba, Ryo Yoshimura, Kyoko Yasuda, Keiko Iida, Naomi Ohsumi, Daniel Sauter

**Hokkaido University, Japan**
Takasuke Fukuhara, Tomokazu Tamura, Rigel Suzuki, Saori Suzuki, Hayato Ito, Keita Matsuno, Hirofumi Sawa, Naganori Nao, Shinya Tanaka, Masumi Tsuda, Lei Wang, Yoshikata Oda, Marie Kato, Zannatul Ferdous, Hiromi Mouri, Kenji Shishido

**Tokyo Metropolitan Institute of Public Health, Japan**
Kenji Sadamasu, Kazuhisa Yoshimura, Hiroyuki Asakura, Isao Yoshida, Mami Nagashima

**Tokai University, Japan**
So Nakagawa, Jiaqi Wu

**Kyoto University, Japan**
Kotaro Shirakawa, Akifumi Takaori-Kondo, Kayoko Nagata, Yasuhiro Kazuma, Ryosuke Nomura, Yoshihito Horisawa, Yusuke Tashiro, Yugo Kawai, Kazuo Takayama, Rina Hashimoto, Sayaka Deguchi, Yukio Watanabe, Ayaka Sakamoto, Naoko Yasuhara, Takao Hashiguchi, Tateki Suzuki, Kanako Kimura, Jiei Sasaki, Yukari Nakajima, Hisano Yajima

**Hiroshima University, Japan**
Takashi Irie, Ryoko Kawabata

**Kyushu University, Japan**
Kaori Tabata

**Kumamoto University, Japan**
Terumasa Ikeda, Hesham Nasser, Ryo Shimizu, MST Monira Begum, Otowa Takahashi, Kimiko Ichihara, Takamasa Ueno, Chihiro Motozono, Mako Toyoda

**University of Miyazaki, Japan**
Akatsuki Saito, Maya Shofa, Yuki Shibatani, Tomoko Nishiuchi

**Acknowledgments**

We would like to thank all members of The Genotype to Phenotype Japan (G2P-Japan) Consortium. We thank Dr. Kenzo Tokunaga (National Institute of Infectious Diseases, Japan) for sharing materials and Dr. Jin Kuramochi (Interpark Kuramochi Clinic, Japan) for sharing human sera. We gratefully acknowledge the numerous laboratories worldwide that have provided sequence data and metadata to GISAID. A full list of originating and submitting laboratories for the sequences used in our analysis can be found at https://www.gisaid.org using the EPI-SET-ID: EPI_SET_230113qo.
